# Quantitative Receptor Model for Responses That Are Left- or Right-Shifted Versus Occupancy (Are More or Less Concentration Sensitive): The SABRE Approach

**DOI:** 10.1101/2023.08.02.551700

**Authors:** Peter Buchwald

## Abstract

Simple one-to three-parameter models routinely used to fit typical dose-response curves and calculate EC_50_ values using the Hill or Clark equation cannot provide the full picture connecting measured response to receptor occupancy, which can be quite complex due to the interplay between partial agonism and (pathway-dependent) signal amplification. The recently introduced SABRE quantitative receptor model is the first one that explicitly includes a parameter for signal amplification (*γ*) in addition to those for binding affinity (*K*_d_), receptor-activation efficacy (*ε*), constitutive activity (*ε*_R0_), and steepness of response (Hill slope, *n*). It can provide a unified framework to fit complex cases, where fractional response and occupancy do not match, as well as simple ones, where parameters constrained to specific values can be used (e.g., *ε*_R0_=0, *γ*=1, or *n*=1). Here, it is shown that SABRE can fit not only typical cases where response curves are left-shifted compared to occupancy (*κ*=*K*_d_/EC_50_>1) due to signal amplification (*γ*>1), but also less common ones where they are right-shifted (i.e., less concentration-sensitive; *κ*=*K*_d_/EC_50_<1) by modeling them as apparent signal attenuation/loss (*γ*<1). Illustrations are provided with *μ*-opioid receptor (MOPr) data from three different experiments with one left- and one right-shifted response (G protein activation and *β*-arrestin2 recruitment, respectively; EC_50,Gprt_<*K*_d_<EC_50,βArr_). For such cases of diverging pathways with differently shifted responses, partial agonists can cause very weak responses in the less concentration-sensitive pathway without having to be biased ligands due to the combination of low ligand efficacy and signal attenuation/loss – an illustration with SABRE-fitted oliceridine data is included.

## Introduction

### Quantitative receptor models and their parametrization

Receptors [1, 2] are at the core of our current understanding of mechanism of drug action [3-5]. By now, it is also well established that the relationship between receptor occupancy and response can be quite complex. Being able to connect the concentration of the agonist or antagonist ligand to the response it causes, i.e., establishing concentration-response relationships, is of major interest in general and a main goal in quantitative pharmacology. It is now clear that to be able to do so one needs to consider not just *(i)* the degree of receptor occupancy (binding) but also that *(ii)* an agonist occupied receptor is not necessarily active (partial agonism), *(iii)* an unoccupied receptor is not necessarily inactive (constitutive activity), *(iv)* full or close to full responses can be produced even when only a fraction of receptors is occupied and/or active (signal amplification creating the appearance of “spare receptors” or “receptor reserve”), and *(v)* changes in ligand concentration may produce more or less abrupt changes in response than predicted by a classic law of mass action. Therefore, a quantitative receptor model that can account for all these needs some parametrization to characterize:

i. the ability of the ligand to bind the receptor (*affinity*),
ii. the ability of the ligand to activate the receptor upon binding (*efficacy*) – something that has been known since at least the mid-1950s following the work of Ariëns [6] and Stephenson [7] and is the basis of minimal two-state models,
iii. the degree of activation of unoccupied receptors (*efficacy of constitutive activity*) – as it became clear with the realization that there are constitutively active receptors in the 1980s (see [8, 9]) and led to the need for full two-state models,
iv. the possibility of pathway-dependent signal amplification that causes concentration–response curves to be shifted to the left compared to concentration-occupancy curves (*amplification* or *gain*), and
v. the steepness of concentration dependence (*Hill slope* or *coefficient*) since responses as a function of concentration can change more (or less) abruptly than predicted by a straightforward law of mass action.

Thus, a quantitative model that could account for all these should have at least five parameters. Ideally, assuming that these are independent (at least to a reasonable degree), the model should also be reducible to simplified versions by constraining each of its parameters to fixed values if the corresponding phenomenon is not relevant. For example, if there is no constitutive activity, the corresponding efficacy parameter should be null. Along similar lines, if there is no need for altering the abruptness of the response function from that predicted by a straightforward law of mass action, a Hill slope of unity can be used (*n* = 1) reducing the well-known Hill equation to the simpler and more widely used Clark equation:

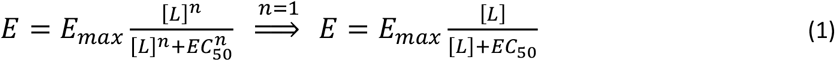

Here, we will use such equations in a normalized (fractional response) form as shown below, i.e., normalized to the *E*_max_ of the assay with *e*_max_ (0 < *e*_max_ ≤ 100%) being the maximum achievable for a given agonist (0 < *f*_resp_ ≤ *e*_max_):

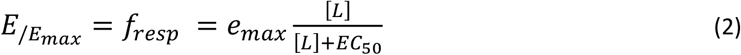

Such two-parameter equations (EC_50_, *e*_max_) are routinely used to fit typical concentration- or dose-response curves – together with three-parameter ones if Hill extensions are allowed (EC_50_, *e*_max_, *n*), and the obtained EC_50_ and *e*_max_ values are considered as indicators of potency and (maximal) efficacy, respectively [10]. Nevertheless, it is also well-recognized that they are not true indicators of the binding affinity (i.e., *K*_obs_ = EC_50_ is not *K*_d_) or the “intrinsic” efficacy of the ligand (i.e., *e*_max_ is no true *efficacy* as even known weak partial agonists could produce full or close to full responses is some systems).

Common quantitative receptor models used to fit more complex cases typically rely on the so-called operational (Black & Leff) model [11], which has a mathematical form identical to that of the minimal two-state (del Castillo-Katz) model [12] and employs one affinity- (*K*_D_) and one efficacy-type parameter (*τ*) [3]:

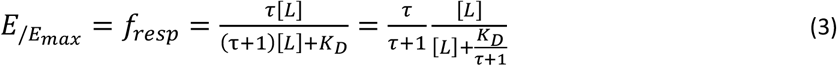

These models, however, use parameters that are nonintuitive, difficult to interpret (i.e., lack clear pharmacological meaning), and still no true indicators of binding affinity (i.e., *K*_D_ is not *K*_d_ and experimental *K*_d_s cannot be used as *K*_D_) or “intrinsic” efficacy (i.e., the *τ* “transducer ratio” is not a true efficacy, and full or close to full agonists need infinitely large *τ*). These models are also cumbersome and difficult to fit in well-defined manner, as it has been highlighted in several papers [13-17]. If just functional data are available (i.e., single concentration-response curves), only the so-called “transduction coefficient” *τ*/*K*_D_ and not *K*_D_ and *τ* independently can be estimated due to identifiability issues during regression [13-17]. Nevertheless, several further variations of this operational model-based equation have been proposed including some with additions needed for constitutive activity [18-24].

Here, after a brief review of the recently introduced SABRE quantitative receptor model [25, 26], the connection between receptor occupancy and response data will be discussed including for less common cases where responses are not left-but right-shifted compared to occupancy (implying log EC_50_ > log *K*_d_). It will be shown that they can be fitted with SABRE by simply allowing its amplification parameter to be less then unity (i.e., as an apparent signal attenuation/loss), and illustrations will be provided with *μ*-opioid receptor (MOPr) data from three experiments where from two responses initiating from the same receptor, one is left- and one is right-shifted compared to occupancy (G protein activation and *β*-arrestin2 recruitment, respectively; EC_50,Gprt_ < *K*_d_ < EC_50,*β*Arr_).

## Methods

### Experimental Data

Experimental data used here are from three different published works performed in three different laboratories that, however, all measured two different activities as well as receptor binding for the *μ*-opioid receptor (MOPr) and all included DAMGO (D-Ala^2^, N-MePhe^4^, Gly-ol^5^–enkephalin) and morphine as agonist: (1) J. McPherson and coworkers from the University of Bristol (Bristol, UK) with HEK293 cells stably expressing T7-tagged MOPr (used to measure agonist-induced [^35^S]GTP*γ*S binding and arrestin-3 (*β*-arrestin2) recruitment using the DiscoveRx PathHunter (plus binding experiments with [^3^H]naloxone) [27]; (2) J. D. Hothersall and coworkers from Pfizer (Cambridge, UK) with HEK293 cells expressing MOPr^wt^ quantifying agonist G_*α*i/o_ responses using a femto HTRF cAMP assay and *β*-arrestin2 recruitment using the DiscoverX PathHunter assay (plus binding via a radioligand binding assay measuring the displacement of [^3^H]diprenorphine) [28]; and (3) M. F. Pedersen and coworkers from the University of Copenhagen (Copenhagen, Denmark) with HEK293A cells using BRET assays to measure G_*α*i2_ activation as well as *β*-arrestin2 recruitment (plus binding measurements with [^3^H]naloxone) [29]. In all three, binding affinities (equilibrium dissociation constants, *K*_d_) were calculated using the Cheng-Prusoff equation [30] (or equivalent corrections in GraphPad Prism) to account for radioligand concentration. Pharmacodynamic parameters used here for DAMGO and morphine (EC_50_, *E*_max_; Table 1) are from these works as published; data used for fitting were obtained from the figures using WebPlotDigitizer [31].

**Table 1.** Summary of the experimental receptor binding and activity data used, and the corresponding fit parameters obtained in the present work.

| Study | McPherson [27] |  | Hothersall [28] |  | Pedersen [29] |  |
| --- | --- | --- | --- | --- | --- | --- |
| Param. \ Cpd. | DAMGO | Morphine | DAMGO | Morphine | DAMGO | Morphine |
|  | **Experimental <sup>a</sup>** |  |  |  |  |  |
| $\log K_d$ | -6.64 | -6.60 | -6.62 | -6.76 | -7.34 | -7.02 |
| $\log EC_{50, Gprt}$ | -7.95 | -7.01 | -7.72 | -7.24 | -8.61 | -8.17 |
| $\log EC_{50, \beta Arr}$ | -6.38 | -6.49 | -6.02 | -6.54 | -6.10 | -6.00 |
| $E_{max, Gprt}$ | 100.0 | 94.2 | 96.8 | 92.9 | 99.0 | 98.0 |
| $E_{max, \beta Arr}$ | 99.2 | 15.2 | 77.7 | 7.5 | 99.0 | 25.0 |
| $\kappa_{Gprt}$ | 20.36 | 2.56 | 12.59 | 3.02 | 18.62 | 14.13 |
| $\kappa_{\beta Arr}$ | 0.551 | 0.776 | 0.251 | 0.603 | 0.058 | 0.095 |
|  | **SABRE <sup>b</sup>** |  |  |  |  |  |
| $\gamma_{Gprt}$ | 30.84 | | 9.88 | | 15.51 | |
| $\gamma_{\beta Arr}$ | 0.669 | | 0.193 | | 0.059 | |
| $\varepsilon$ | 1.000 | 0.178 | 0.948 | 0.375 | 0.999 | 0.823 |
| $\kappa_{Gprt}$ | 30.84 | 6.31 | 9.41 | 4.33 | 15.49 | 12.94 |
| $\kappa_{\beta Arr}$ | 0.669 | 0.941 | 0.235 | 0.698 | 0.06 | 0.226 |

<sup>a</sup> Experimental data ( $\log K_d$ , $\log EC_{50}$ ) are directly from the corresponding publications with $\kappa$ (fold shifts vs occupancy) calculated from the $K_d/EC_{50}$ values (eq. 13). <sup>b</sup> SABRE parameters ( $\gamma$ , $\varepsilon$ ) are those obtained from fitting the data in GraphPad Prism with $\kappa$ calculated using eq. 15.

### Implementation and Data Fitting

All data used here are normalized and have no baseline (i.e., they are in the 0–100% range) and were fitted using GraphPad Prism (GraphPad, La Jolla, CA, USA, RRID:SCR_002798). Fittings with SABRE were done with a custom implementation corresponding to the general eq. 5, which was made available for download (see [26]), and with parameters constrained as indicated for each case.

## Results and Discussion

### SABRE – a quantitative receptor model incorporating signal amplification

The recently introduced SABRE model is the first quantitative receptor model that explicitly includes parametrization for signal amplification via a dedicated *γ* (gain) parameter as implied by its acronym (Signal Amplification, Binding affinity, and Receptor-activation Efficacy) [25, 26]. Its simplified three-parameter version uses *K*_d_ for binding affinity, *ε* for efficacy, and *γ* for signal amplification:

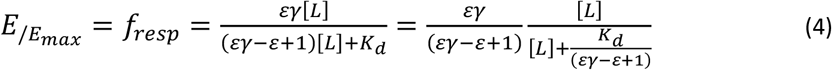

In line with the ideas highlighted in the Introduction, the full general form of SABRE employs the full spectrum of five parameters mentioned there *(i–v)* by also incorporating a Hill coefficient (*n*) and a parameter for constitutive activity (*ε*_R0_) [25, 26]:

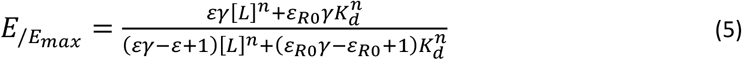

For the sake of simplicity, these two parameters will be considered as fixed (*n* = 1 and *ε*_R0_ = 0) and only the resulting simplified form of SABRE shown in eq. 4 will be used from here on. As highlighted by the rightmost form of this equation (eq. 4), SABRE corresponds to a classic hyperbolic relationship between (fractional) response, *E*/*E*_max_, and ligand concentration, [L], which is sigmoid on the semi-log scale typically used, *E*_/*E max*_ *= ℱ(log[L])*, with an apparent *K*_obs_ (EC_50_) and *e*_max_ (0 < *e*_max_ ≤ 100%) that are:

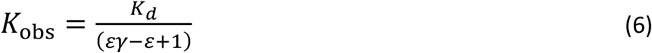

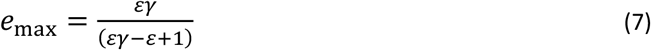

Thus, if only (baseline corrected) concentration-response data are available, the most common case, a constrained form of SABRE with only two adjustable parameters, *K*_d_ and *ε*, can be used for fitting with classic hyperbolic response functions. Constraining *γ* = 1 as a fixed parameter (no amplification), results in *K*_obs_ = *K*_d_ and *e*_max_ = *ε* from equations 6 and 7 above and a corresponding simple Clark-type equation:

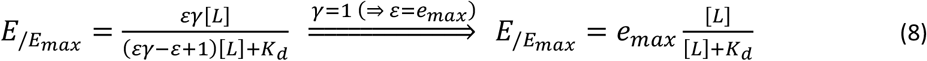

If additional data are available, e.g., multiple test compounds with experimental binding affinities (log *K*_d_) also measured in the same setup, SABRE can provide insight into the signal amplification of the assayed pathway (*γ*) and the efficacies of the test compounds (*ε*) [25, 26, 32]. Since SABRE is the first model that uses explicit parametrization for (post-receptor) signal modulation, it allows a better separation of the receptor binding, receptor activation, and signal transduction (amplification) steps, which can be characterized and quantified via their own distinct parameters: *K*_d_, *ε*, and *γ*, respectively. Thus, SABRE makes possible a clearer conceptualization of receptor signaling [25, 26, 32] with parameters that are more intuitive and easier to interpret than those of the operational model or minimal two-state model (*K*_D_, *τ*; eq. 3). For the same reason, SABRE can fit both simple and complex cases with the same equation (eq. 5) since it can be collapsed into consecutive simplified forms by fixing its parameters at special values as described before (e.g., *ε*_R0_ = 0 if there is no constitutive activity; *n* = 1 if there is no need for non-unity Hill slope, i.e., standard law of mass action responses only; *γ* = 1 if there is no amplification – or if it cannot be reliably evaluated due to lack of enough data; and *ε* = 1 if there is no partial agonism; see [26] for more detail).

### Connecting receptor response and occupancy data

Even if most pharmacological works focus solely on fitting response data only to establish pharmacodynamic (PD) parameters (e.g., EC_50_), a mechanistically relevant receptor model should be able to connect ligand concentration to resulting receptor occupancy and caused response(s). However, these relationships can be complex, and receptor response and occupancy rarely overlap as in Figure 1A. Typically, they are shifted (Figure 1B) for the reasons mentioned in the Introduction that include receptor activation efficacy and signal amplification, which can vary depending on the pathways or vantage points used for assessment. Figure 1B shows an illustrative example with two different responses obtained from the same occupancy, e.g., generated by two pathways with different amplifications including the possibility that the ligand has different abilities (efficacies) to activate these pathways. Here, the typical hyperbolic concentration-dependency of the responses and occupancies as well as the corresponding response versus occupancy curves are shown (top and bottom rows, respectively; being sigmoid on the semi-log scales used here). As both response curves (blue) are left-shifted compared to occupancy (green in Figure 1B, top; *K*_d_ > *K*_obs_ = EC_50_) due to signal amplification, responses run ahead of occupancy (except for the rightmost part of response 2) as more clearly evident in the curves of the bottom response vs occupancy graph of the same Figure 1B. *Formalism linking classic hyperbolic functions (with Κ shift parameter)*

**Figure 1.**
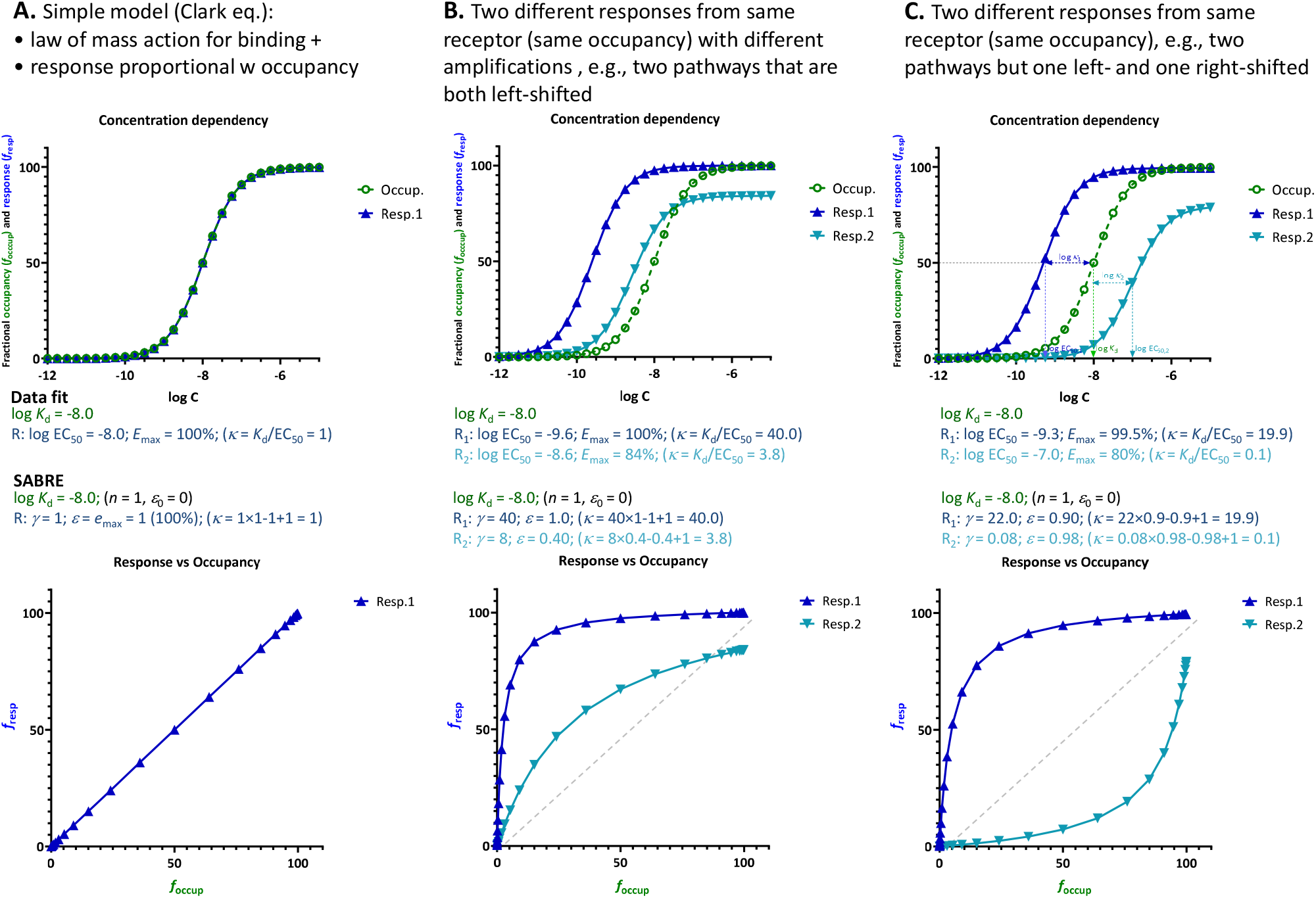
Illustrations of the relationship between fractional response (*f*_resp_=*E*/*E*_max_) and occupancy (*f*_occup_ = [LR_occup_]/[LR_max_]) for (**A**) the simplest case corresponding to the Clark equation (i.e., law of mass action for binding and receptor response proportional with number of occupied receptors), (**B**) two different responses obtained from the same receptor (same occupancy), e.g., two pathways with different amplifications resulting in two left-shifted responses as compared to the occupancy, and (**C**) two different responses obtained from the same receptor (same occupancy) but one left- and one right-shifted as compared to the occupancy. Top row: receptor responses (blue) and occupancies (green) as a function of ligand concentration on typical semi-log scales (i.e., shown as a function of log *C* = log [L]). Bottom row: corresponding response versus occupancy curves (*f*_resp_ as a function of *f*_occup_). Parameter values shown in the middle are those obtained when fitting these data with separate sigmoidal equations (such as eq. 1, which are commonly used but cannot connect *K*_d_, EC_50_, and *E*_max_ values) and by SABRE (eq. 4) using a fixed *K*_d_ (as defined by the binding data) and gain (*γ*) and efficacy (*ε*) parameters as indicated. In all cases, the values of *Κ* (i.e., fold change in response vs occupancy at the half-maximal values, eq. 13) are also indicated in parentheses. While these are simulated data generated with SABRE, experimental data corresponding cases like these are available in the literature (see, e.g., Figure 2 and Figure 3).

Assuming that (fractional) occupancy and response are described by classic hyperbolic functions versus ligand concentration [L] (sigmoid on log-scale), they can be written in the general form of eq. 2 as:

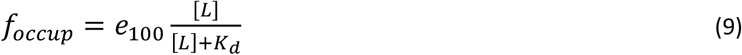

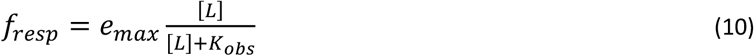

Occupancy is assumed to always reach a maximum of 100% (hence, the *e*_100_ notation), while responses can plateau at smaller values for partial agonists (0 < *e*_max_ ≤ 100%). To connect *f*_resp_ and *f*_occup_ and obtain the general functional form illustrated in the bottom row of Figure 1 (essentially the general form connecting two different hyperbolic functions of ligand concentration characterized by *e*_max_ & *K*_obs_ and *e*_100_ & *K*_d_, respectively), one can express [L] from eq. 9

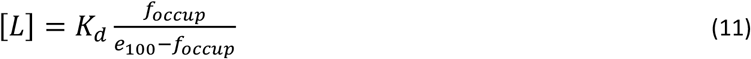

and plug it in eq. 10, giving

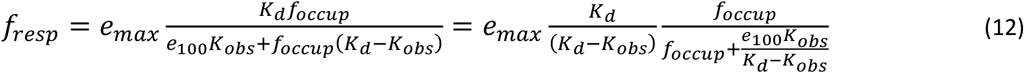

Thus, response (*f*_resp_, characterized by *e*_max_ and *K*_obs_ = EC_50_) is connected to occupancy, (*f*_occup_, characterized by *e*_100_ = 1 and *K*_d_) via a hyperbolic function. Introducing *Κ* as a parameter quantifying the shift between response and occupancy (essentially a fold change in response vs occupancy at the half-maximal values)

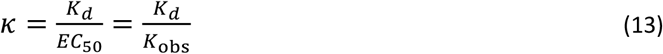

*f*_resp_ can be written as:

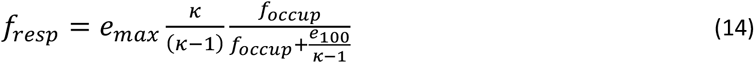

*Formalism in SABRE*

Notably, these general functional dependencies that rely only on the assumption of hyperbolic (sigmoid on log scale) functions (eq. 9, 10, and 14; Figure 1) fit well within the formalism of SABRE. The concentration-response form of SABRE (eq. 4) links directly to the general sigmoid response function (eq. 10, identical with eq. 2) via equations 6 and 7 that define *K*_obs_ and *e*_max_ in terms of the *ε* and *γ* parameters of SABRE. Equations 6 and 7 can also be used to connect *f*_resp_ to *f*_occup_ as in eq. 14 above just using the parameters of the SABRE model. From eq. 6 for *K*_obs_,

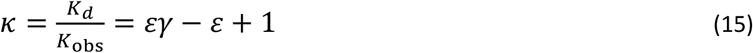

Thus

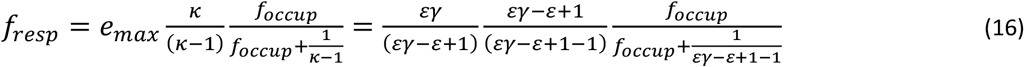

giving the same form that has been deduced before [25, 26]:

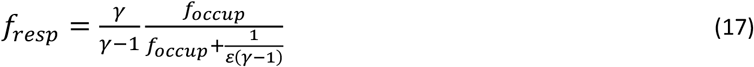

Hence, SABRE can fit either the hyperbolic concentration-response functions directly using eq. 4 or the corresponding (also hyperbolic) response versus occupancy functions using eq. 17, and the results should be the same; an example is included below (Figure 2, Supplementary information, Table S1). When fitting with SABRE, there can be some ambiguity if there are not enough and sufficiently wide-spread data, especially as the efficacy values are dependent on the common gain parameter, and it is not always evident what is a true full agonist in the assay (except that the maximum efficacy is limited at *ε* = 1.0). Nevertheless, if there are enough good quality data covering an adequately large range, well-defined fit can be obtained.

**Figure 2.**
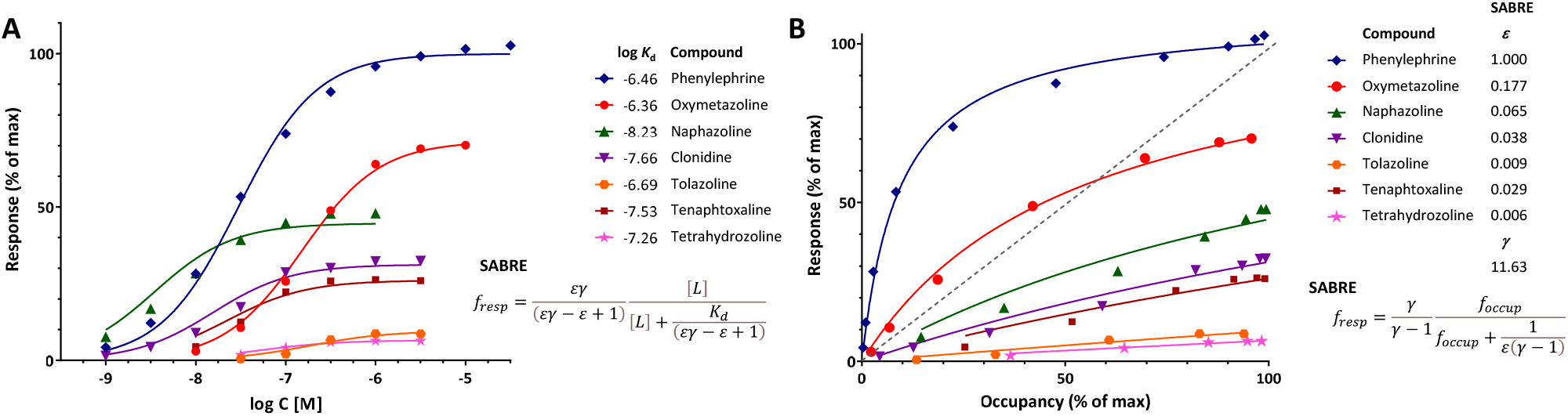
Illustration of fit of complex concentration-response data with SABRE. Data are for a series of imidazoline-type *α*-adrenoceptor agonists for which both response and binding were measured [33]. (**A**) Concentration-response data of seven compounds (symbols) fitted with SABRE using the equation shown (eq. 4) with experimental log *K*_d_ values as shown and only eight adjustable parameters: a common amplification (gain, *γ*) and seven individual efficacies (*ε*). (**B**) Normalized response versus occupancy data for the same compounds (symbols) and their corresponding fit with SABRE using the equation shown (eq. 17) and the same parameters as above. Due to the interplay between amplification and partial agonism, fractional (normalized) response can either exceed or lag occupancy, as comparison to the dashed unity line in the bottom figure clearly reveals.

#### Fit of response and occupancy data for α-adrenergic agonists

An illustration of fit with SABRE using receptor binding (*K*_d_) data assessed together with response is provided with the concentration-dependent contractions of isolated rat aorta induced by a series of imidazoline type *α*-adrenoceptor agonists such as phenylephrine, oxymetazoline, naphazoline, clonidine, tolazoline, and others [33]. Response data for seven compounds, which requires 14 parameters to fit with standard sigmoid curves (7 compounds × 2 parameters, EC50 and *e*_max_ for each), can be fitted with SABRE without significant loss in the quality of fit (Figure 2) while using only 8 parameters (1*γ* + 7*ε*s) and also integrating the experimental *K*_d_s (Supplementary information, Table S1). Thus, SABRE, the more parsimonious model, gives a fit that is only slightly worse (as judged based on the sum of squared errors, SSE, which increased from 91.2 to 211.2, and the corrected Akaike Information Content, AICc, which increased from 81.2 to 93.9) while it also provides valuable additional insight, as it integrates the binding data, suggests a value of the amplification along the assayed pathway (*γ* = 11.6 ± 1.8), and quantifies the efficacies of the evaluated agonists (*ε* ranging from 1.0 for phenylephrine to 0.006 for tetrahydrozoline).

The shift between response and occupancy, as quantified by *Κ* (eq. 13) from the experimental data for the full antagonist phenylephrine, provides context for the gain parameter *γ* of SABRE (and its interplay with *ε*). As shown in eq. 15, *Κ* can be written as *εγ* – *ε* +1 in terms of SABRE parameters, so that for a full agonist (*ε* = 1), *Κ* = *γ*. Here, *γ* estimated from fitting the whole dataset (11.63 ± 1.83) indeed agrees very well with *Κ* of the full agonist phenylephrine (12.3) (Table S1). For the other compounds, which are all partial agonists, the shifts are smaller, but they are also well reproduced by the SABRE estimates (Table S1) with the note that the EC_50_ and, hence, *Κ* estimates for the weak partial agonists (e.g., those with *e*_max_ < 30%) cannot be considered reliable as well-defined values could not be obtained for them due to the limited range of responses.

### Right-shifted response data

Cases where the response is not left-but right-shifted compared to occupancy (implying *Κ* < 1 as EC_50_ > *K*_d_, see Figure 1C) are less common but have been documented. Illustrative examples obtained in three different works with the *μ*-opioid receptor (MOPr) involving *β*-arrestin recruitment are shown in Figure 3. Importantly, these right-shifted responses are one of two responses assessed that are generated along different pathways originating from the same receptor with the other one, G-protein activation, being left-shifted. Pathways are defined by a transducer protein or family thereof, binding intracellularly to the receptor and eliciting a distinct cellular downstream signaling cascade, trafficking, or internalization as per current IUPHAR guidelines [34]. Some receptors can engage multiple downstream signaling pathways (i.e., are pleiotropically linked), can activate them differentially, and can do so in a tissue-dependent manner. For example, for G-protein coupled receptors (GPCRs), according to present knowledge, this includes four G protein families (the G_s_, G_i/o_, G_q/11_, and G_12/13_ pathways) and the GPCR kinase (GRK) and arrestin families (total of six transducer protein families) [34].

**Figure 3.**
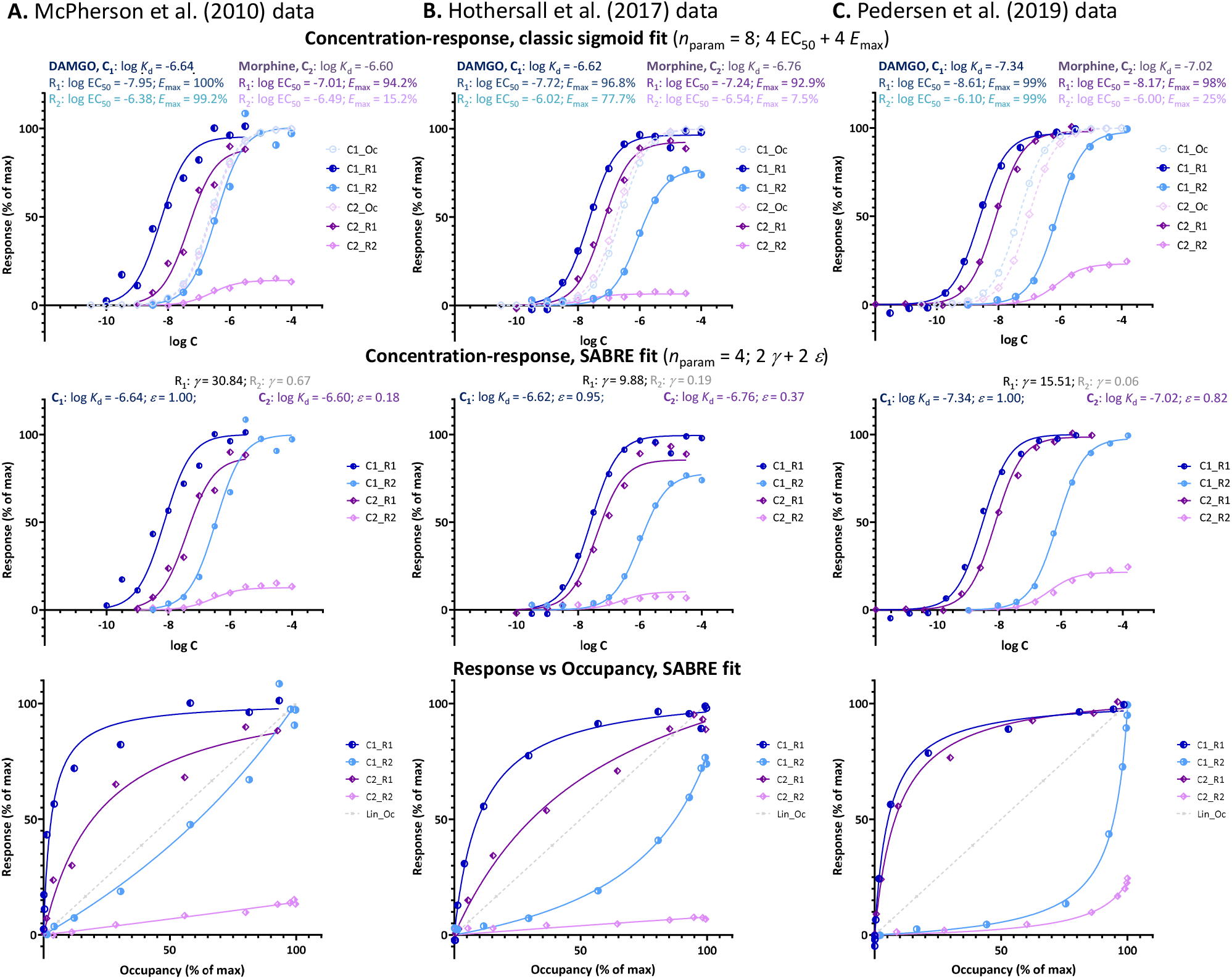
Fit of DAMGO and morphine induced *μ*-opioid receptor (MOPr) responses along two different pathways involving G-protein activation and *β*-arrestin2 recruitment, respectively as obtained in three different works by three different groups: (**A**) McPherson et al. at the University of Bristol [27], (**B**) Hothersall et al. at Pfizer [28], and (**C**) Pedersen et al. at the University of Copenhagen [29]. Data and fit for DAMGO (C_1_) and morphine (C_2_) shown in blue and purple, respectively with G-protein activation (R_1_) in darker and *β*-arrestin recruitment responses (R_2_) in lighter shades. Top row: fit with classic sigmoid concentration-response curves and corresponding binding affinity (*K*_d_) and pharmacodynamic (EC_50_, *E*_max_) parameters as obtained in these works (with calculated occupancy curves shown as dashed lines). Middle row: fit of the same data with SABRE using only four parameters for each dataset – two pathway amplifications (*γ*) and two ligand efficacies (*ε*). Bottom row: corresponding response versus occupancy graphs; the stronger the curvature here compared to the straight unity line of linear response, the more the left- or right-shift in the concentration-response curve as compared to the occupancy one.

Such right-shifted responses, which are a main focus of the present work, are usually considered indications of an occupancy threshold issue, i.e., indications that the receptor concentrations are not negligible compared to those of the ligand – an otherwise common assumption in pharmacology [35]. However, in cases such as those shown in Figure 3, this is unlikely since, as mentioned, they are one of two responses generated at the same receptor (here, MOPr) along different pathways with one (G protein activation) being clearly left-shifted and the other (*β*-arrestin2 recruitment) clearly right-shifted in all three experiments (EC_50,Gprt_ < *K*_d_ < EC_50,*β*Arr_ so that *Κ*_Gprt_ > 1 and *Κ*_*β*Arr_ < 1; Table 1). Notably, for the cases shown here (Figure 3), both the left- and right-shifted responses seem to maintain the classic hyperbolic shape (sigmoid on log-scale).

Left-shifted responses, which are more concentration-sensitive than occupancy, typically result from some type of signal amplification with the response plateauing at a maximum due to reaching limiting saturation in the final step. This is most clearly evident in response versus occupancy figures, such as those shown in the bottom rows of Figure 1 and Figure 3, where response is running “ahead” of occupancy, *f*_resp_ > *f*_occup_ (dark blue vs dotted gray line). For example, for Resp.1 in Figure 1C, a 20% (fractional) occupancy, *f*_occup_ = 20%, already results in close to maximum response, *f*_resp1_ ≈ 90%. There are many similar or even more extreme cases documented [25] such as, e.g., • the response of human calcitonin receptor type 2 to calcitonin, where *f*_occup_ = 20% produces almost full response (*f*_resp_ ≈ 100%) [36], • the response of guinea pig ileum to histamine, where *f*_occup_ as low as 2% already produces almost full response [5, 37, 38], or • the stimulation of *β*-adrenergic receptors in the heart by epinephrine, where *f*_occup_ = 1–3% in rats and *f*_occup_ = 10–20% in humans produces half-maximal response (*f*_resp_ ≈ 50%) [39]. Also, the G-protein activation response produced by DAMGO at MOPr, where *f*_occup_ = 20% produces close to full response (*f*_resp_ > 80%) as shown in Figure 3. Right-shifted responses such as those shown in Figure 1 and Figure 3 seem to suggest the opposite: an apparent signal attenuation or loss with the response running “behind” the occupancy (*f*_resp_ < *f*_occup_) and, thus, being less concentration sensitive (*Κ* < 1). For example, for Resp.2 in Figure 1C, 80% (fractional) occupancy, *f*_occup_ = 80%, only results in ∼15% of the maximum response, *f*_resp2_ = 15%. For the *β*-arrestin response produced by the full agonist DAMGO at MOPr [29], *f*_occup_ = 75% only produces *f*_resp_ < 15% as shown in Figure 3C.

#### Formalism with hyperbolic functions and in SABRE

As the hyperbolic concentration-response shapes seem to be maintained for the right-shifted cases, the derived equations linking response to occupancy can still be used, just with *Κ* < 1 for eq. 14 (i.e., fold decrease and not increase in response vs occupancy at the half-maximal values). Similarly, right-shifted responses can very nicely be accommodated within the formalism of SABRE by allowing the gain parameter to be less than unity (*γ* < 1). With this, the corresponding equations such as equations 4 and 17 can be used without needing any further modifications. As *γ* > 1 is an indication of signal amplification, this suggests the opposite, i.e., an apparent signal attenuation or loss.

#### Right-shifted β-arrestin responses at MPOr

Data from three different experimental works by three different groups studying possible biased agonism at the MOPr (a class A GPCR) are used here for a SABRE-based quantitative analysis of right-shifted responses. MOPr signaling has been in particular focus recently because of the purported possibility of achieving improved analgesia by reducing the unwanted side-effects of opiate therapeutics via biased signaling. Data used here are from three different published works that, however, all *(i)* included both DAMGO (D-Ala^2^, N-MePhe^4^, Gly-ol^5^–enkephalin) and morphine as agonist ligands, *(ii)* quantified both G-protein activation and *β*-arrestin recruitment in cell-based assays, and *(iii)* in addition to responses also measured MOPr binding affinities: works by McPherson and coworkers at the University of Bristol (UK; 2010) [27], Hothersall and coworkers at Pfizer (Cambridge, UK; 2017) [28], and Pedersen and coworkers at the University of Copenhagen (Denmark; 2019) [29] (see Methods for relevant experimental details). As shown in Table 1, there were some differences in the measured binding (log *K*_d_) and activity (log EC_50_) data among these works; nevertheless, in all cases, the G protein activation responses were left-shifted compared to occupancy (*Κ*_Gprt_ = *K*_d_/EC_50,Gprt_ > 1.0), whereas *β*-arrestin recruitment responses were right-shifted (*Κ*_βArr_ = *K*_d_/EC_50,βArr_ < 1.0) despite originating from the same receptors.

Plotting of the data either as classic concentration-response curves (in parallel with occupancies) or as response vs occupancy (Figure 3, top and bottom rows, respectively), clearly shows that the *β*-arrestin responses lag behind the occupancy even for the full agonist DAMGO (light blue versus dashed curves) while the G-protein responses (dark blue) are well ahead. At the lower end of fractional occupancy where, e.g., only a quarter of the receptors is occupied, *f*_occup_ = 25%, *β*-arrestin responses are minimal, *f*_resp,*β*Arr_ < 5%, whereas corresponding G-protein responses are already approaching their maximum, *f*_resp,Gprot_ > 80% (Figure 3B, C). Thus, there has to be something limiting the *β*-arrestin response as compared to receptor occupancy.

Regardless of this, fitting with SABRE produces good results for both responses in all three datasets (Figure 3) and with consistent sets of parameters (Table 1). In all cases, the G-protein activation responses are amplified with the gain parameter *γ* > 1 with values ranging somewhere between 10 and 30. Meanwhile, the *β*-arrestin responses gave *γ* < 1 with values ranging from ∼1/15 to 1/1.5. Allowing different efficacies for the two divergent pathways (as possible sign of biased agonism [25, 26] – see also below), did not result in significantly different values in any of these cases; therefore, all fittings shown here (Figure 3, Table 1) were obtained with a single efficacy for each ligand (i.e., *ε*s restricted to the same value for both pathways, *ε*_Gprt_ = *ε*_*β*Arr_), which also made the fittings much better defined. Results indicated DAMGO to be a full or very close to full agonist (*ε* ≈ 1.0) and morphine a weaker partial agonist in all three data sets (Table 1).

#### Implications for biased versus weak agonism

For pleiotropically linked receptors such as MOPr responses along different pathways could be different even with the same activation signal, often for obvious physiological reasons. For example, G protein activations often assessed through second messenger assays (e.g., measurement of cAMP) tend to be highly amplified, whereas *β*-arrestin complementation ones do not, resulting in different potencies (e.g., EC_50,Gprt_ << EC_50,*β*Arr_). In addition to this phenomenon, which is termed system bias, ligands might show what has been designated as biased agonism (functional selectivity), i.e., activate these pathways to different degrees even if they originate from the same receptor [40-52]. Biased agonism is an intriguing option for improved therapeutic action, and G-protein biased ligands at MOPr have been particularly pursued following the suggestion that they might be less likely to induce unwanted side effects, such as constipation and respiratory depression, than commonly used opioids due to differential engagement of G proteins versus *β*-arrestins. However, quantifying bias is challenging and might not even be achievable in most cases [16, 48, 53]. Visual evaluations can be done best with bias plots that show the response produced in one signaling pathway as a function of the response produced in the other at the same ligand concentration as shown, for example, in Figure 4D.

**Figure 4.**
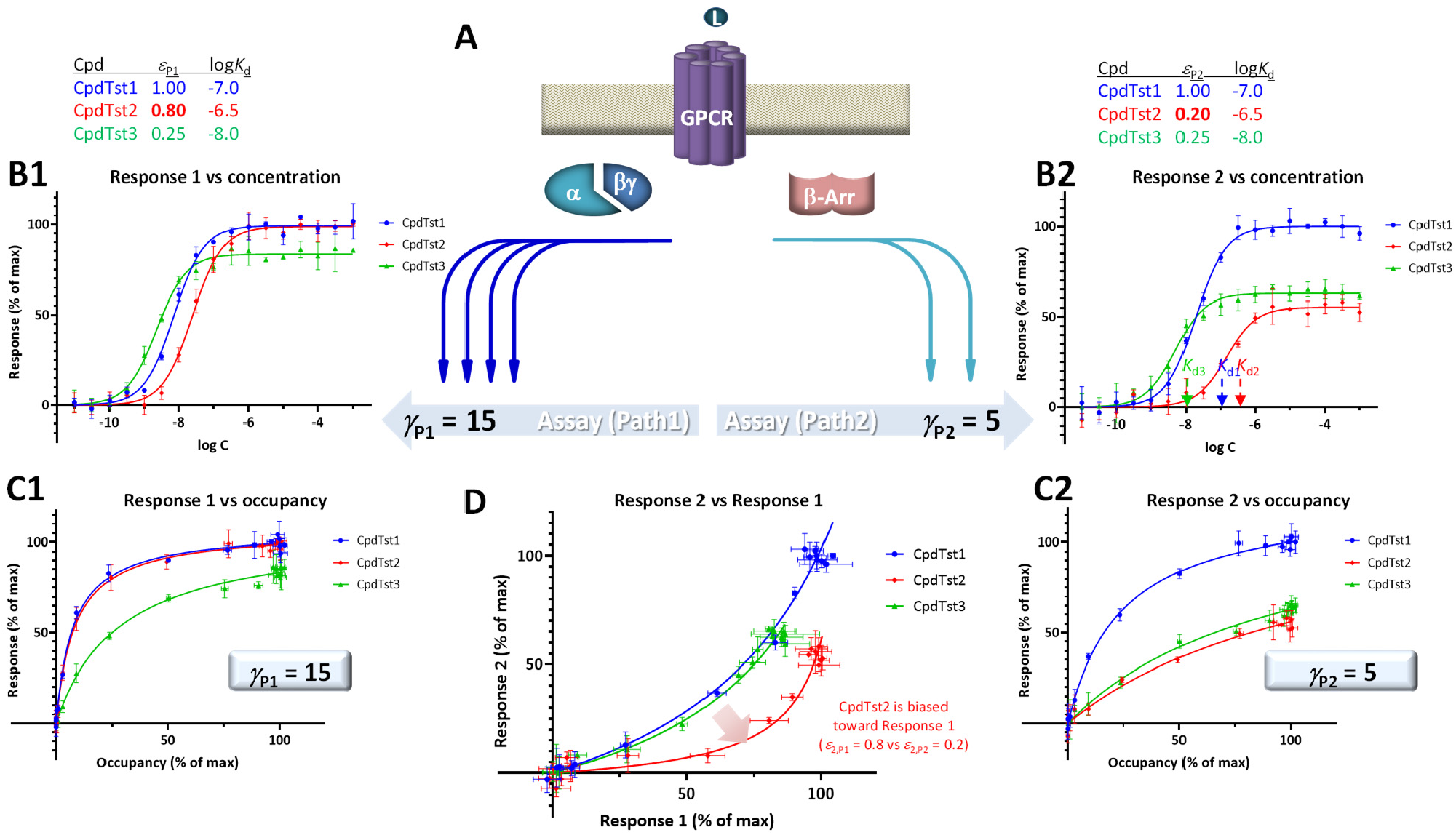
Illustration of biased agonism with response data for two different downstream pathways originating from the same receptor (**A**) generated with the assumption of the present SABRE model. Simulated data (symbols) for three hypothetical compounds were generated using the parameter values shown at top with CpdTst2 (red) having two different efficacies (*ε*_P1_ ≠ *ε*_P2_) as a biased agonist (highlighted in yellow; see text for details). Data for two pathways involving different signal amplifications (*γ*_P1_ = 15 left and *γ*_P2_ = 5 right) are shown as classic semi-log concentration-response curves (*f*_resp_ vs log *C*; **B1, B2**), fractional response versus occupancy curves (*f*_resp_ vs *f*_occup_; **C1, C2**), and a bias plot (*f*_resp1_ vs *f*_resp2_; **D**).

For quantitative assessment, current methods typically rely on calculating ΔΔlog(*τ*/*K*_D_) or ΔΔlog(*E*_max_/EC_50_) versus a selected reference compound [16, 48]. SABRE allows a conceptually different approach as long as there is sufficient data and adequate fit can be achieved [25, 26]. This is done by allowing each ligand to have different, pathway-specific efficacies (not just a single receptor-specific efficacy) and then comparing the obtained fitted values for indication of bias. Efficacy values that are significantly different (*ε*_P__*k*_ ≠ *ε*_P__*l*_) can be considered as indication of biased agonism [25, 26]. If *γ* and *ε* values cannot be obtained in sufficiently well-defined manner, *εγ* products can be compared (with a designated reference compound).

For illustration, a set of data generated within the framework of SABRE for two divergent pathways (P_1_, P_2_) originating from the same receptor but with different amplifications (*γ*_P1_, *γ*_P2_) and assuming three compounds having different affinities (CpdTst1, 2, and 3 with log *K*_d_s of -7.0, -6.5, and -8.0, respectively) is shown in Figure 4. CpdTst1 (blue) was assumed to be a balanced and full agonist for both pathways (*ε*_1,P1_ = *ε*_1,P2_ = 1.0), CpdTst3 (green) a weak balanced agonist (*ε*_3,P1_ = *ε*_3,P2_ = 0.25), and CpdTst2 (red) a biased agonist with higher efficacy for pathway 1 than for 2 (*ε*_2,P1_ = 0.8, *ε*_2,P2_ = 0.2). Corresponding data are shown as typical concentration-response curves (*f*_resp_ vs log *C*; Figure 4B1, B2), response versus occupancy graphs (*f*_resp_ vs *f*_occup_; C1, C2), and classic bias plot used in such cases (*f*_resp1_ vs *f*_resp2_ [43, 54]; D). As the pathways have different amplifications (*γ*_P1_ ≠ *γ*_P2_; here, *γ*_P1_ = 15 and *γ*_P2_ = 5), response plots are curvilinear to different degrees even for balanced (non-biased) ligands (system bias) making it challenging to identify biased agonists using these plots (e.g., Figure 4C1 vs C2). Thus, it is difficult to notice that CpdTst2 is biased (red vs blue and green) except in the bias plot (Figure 4D) directly depicting *f*_resp1_ versus *f*_resp2_. Systems that are strongly biased and have widely different amplifications resulting in highly curved plots can further mask this.

To illustrate the effect of right-shifted responses on this, responses for the same three hypothetical compounds (CpdTsts1–3) but with a second pathways that now has right- and not left-shifted response (*γ*_P2_ < 1) are depicted in Figure 5. Accordingly, the curvature in the corresponding response 2 vs occupancy plot (Figure 5C2 vs Figure 4C2) is different, and because of the signal attenuation, partial agonists produce only weak responses here regardless of whether they are biased or not. With the values used here (*γ*_P1_ = 15 and *γ*_P2_ = 0.15), both compounds 2 and 3 produce only very weak responses in this second pathway; thus, not just biased (CpdTst2, red) but also weak balanced agonists produce no detectable responses (CpdTst3, *ε*_3_ = 0.25, green; Figure 5). Notably, even in the bias plot (Figure 5D) that typically allows the best discrimination, the weak balanced agonist CpdTsts3 groups more with the biased agonist (CpdTsts2) than the balanced full agonists (CpdTst1).

**Figure 5.**
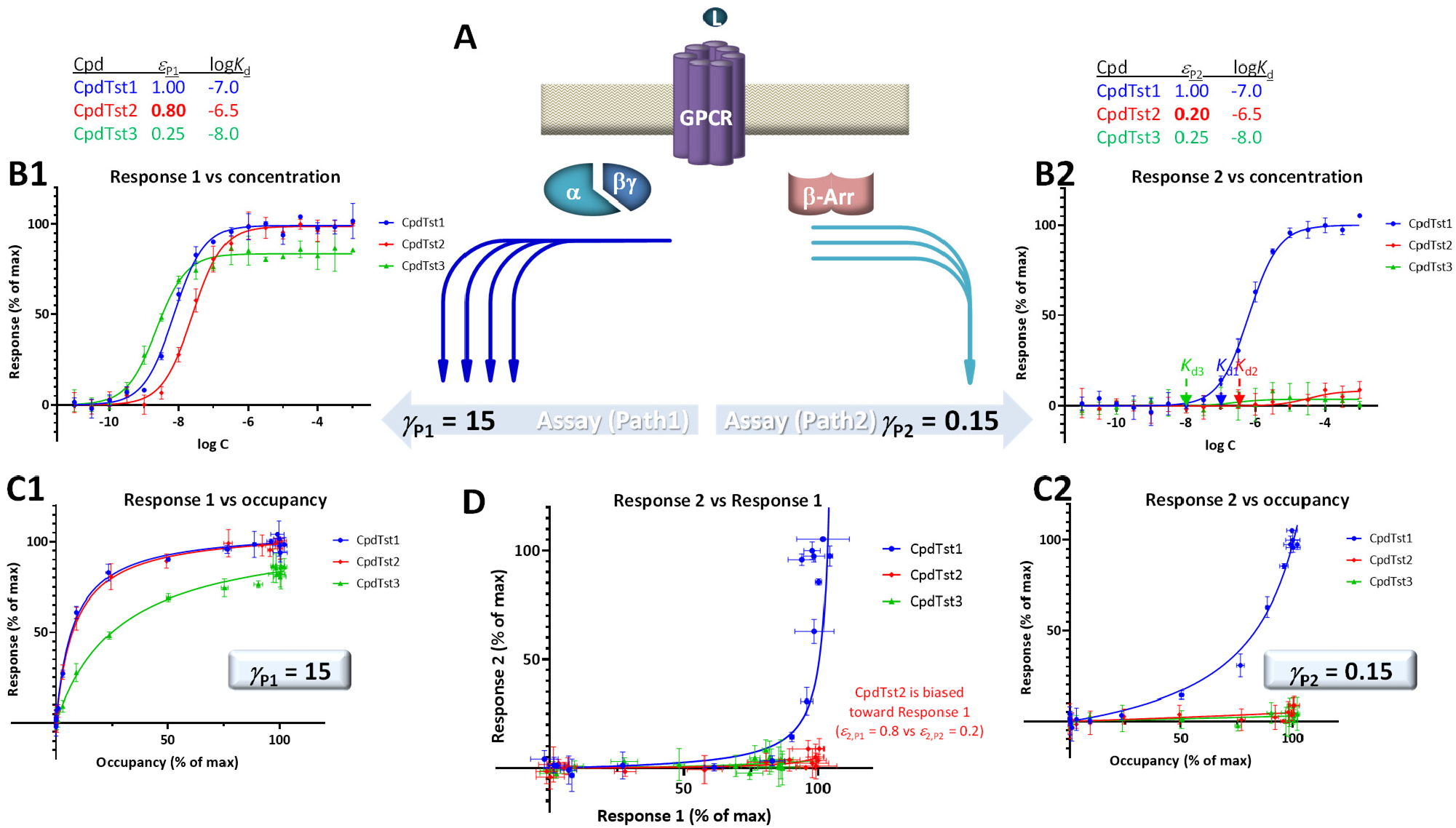
Same as Figure 4 but with response along the second pathways being right-shifted compared to occupancy (*γ*_P2_ = 0.15 < 1.0) to highlight that in this case, even balanced agonist can produce very little activity in this pathway if they are weak partial agonists (CpdTst3). Note that even in the bias plot (D), this weak balanced agonist (green) looks more like the biased agonist (red) than the balanced full agonist (blue).

Such rapid deterioration of responses produced by partial agonists in pathways with right-shifted responses (i.e., with apparent signal attenuation/loss, *γ* < 1) as efficacy decreases is well-illustrated by the *β*-arrestin recruitment data at MOPr as assayed by Pedersen and co-workers [29], part of which was used earlier for DAMGO and morphine (Figure 3C). As for the *β*-arrestin pathway here *γ* << 1, weak agonists such as buprenorphine and oliceridine cause very little response (Figure 6). These data can still be fitted well with SABRE using single efficacies for both pathways (Supplementary information, Table S2); thus, without having to assume biased agonism as discussed before. Note that oliceridine (*R*-TRV130, included here together with its *S* isomer, *S*-TRV130) was developed as a MOPr biased agonist, and it was approved by the FDA for clinical use in 2020 (Olinvyk) as one of the first possible products showing the clinical promise in developing biased agonists [55]. However, it has been suggested that the improved safety profiles of such compounds are due not to the relative reduction in *β*-arrestin mediated signaling because of biased agonism, but to the low intrinsic efficacy in all signaling pathways [56-58]. Fit with SABRE here (Figure 6, Table S2) seems to suggest the same, i.e., that the weak *β*-arrestin recruitment is due to the combination of weak ligand agonism (low *ε*) and signal attenuation / right-shifted response in that pathway (*γ* < 1) without a need for biased agonism.

**Figure 6.**
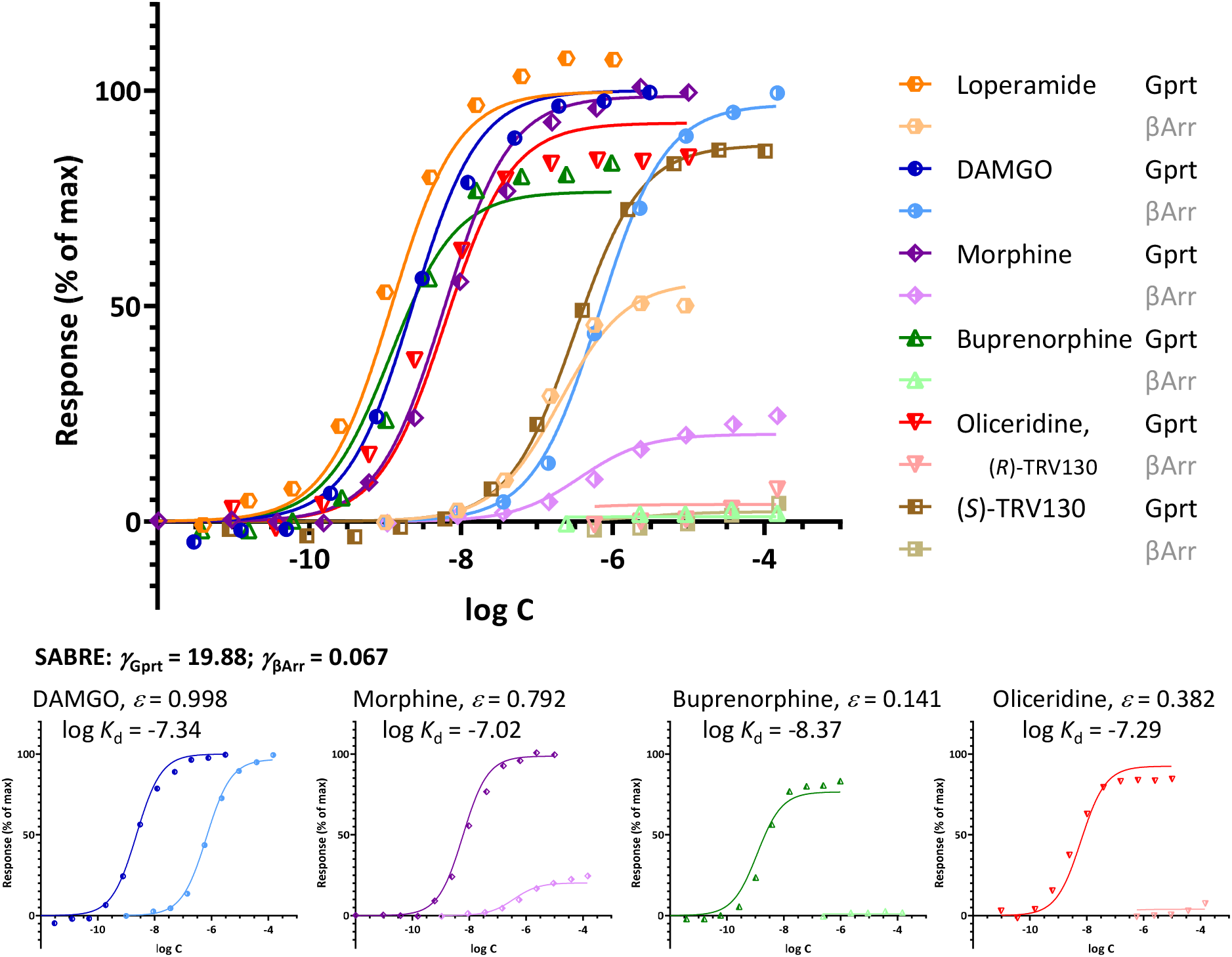
Fit of different agonist-induced MOPr responses along two different pathways involving G-protein activation and *β*-arrestin2 recruitment. Data from [29] are shown as symbols with G-protein activation in darker and *β*-arrestin responses in lighter colors for DAMGO (blue), morphine (purple), buprenorphine (green), oliceridine (red), and its *S* isomer (*S*)-TRV130 (brown). Lines indicate the unified fit obtained with SABRE using experimentally measured log *K*_d_ values, two pathway amplifications (*γ*_Gprt_, *γ*_*β*Arr_), and a single efficacy parameter (*ε*) for each agonist. Due to signal amplification in the G protein response pathway (*γ*_Gprt_ > 1) and apparent attenuation / loss in the *β*-arrestin2 one (*γ*_*β*Arr_ < 1), weak agonists (*ε* << 1.0) such as buprenorphine (*ε* = 0.141) and oliceridine (*ε* = 0.382), produce essentially no *β*-arrestin response even if no bias is assumed (*ε*_Gprt_ = *ε*_*β*Arr_).

#### Mechanistic possibilities

Such right-shifted responses mean that responses are still far from the maximum when occupancy is already approaching saturation (e.g., at [L] = 10×*K*_d_ where *f*_occup_ = 91%) and then catch up abruptly (Figure 1C, bottom). As mentioned, right-shifted responses are usually considered indications of an occupancy threshold issue where receptor concentrations are not negligible compared to ligand concentrations. However, for cases such as those discussed here involving responses generated by the same ligands at the same receptors with one response being left- and one right-shifted (e.g., Figure 3, Figure 6), it is unlikely that the right-shift results from an occupancy threshold issue. Some pathway-specific threshold issue such as one along the *β*-arrestin pathway that does not affect the G protein activation pathway might be a possibility. For example, a need to reach some minimum threshold of activation, including a threshold of sufficiently prolonged activation, or a need for reenforced signals that have to come from multiple occupied receptors within the same system. Loss of weak overall signal or some other issue could also be involved.

Notably, response of the *β*-arrestin pathway for receptors other than MOPr does not appear to be right-shifted (*Κ*_*β*Arr_ < 1.0), at least, for the few cases where occupancy data were also assessed in the same work. For example, for epinephrin at *β*_2_ adrenergic receptors: log *K*_d_ = -6.54, log EC_50,Gprt,cAMP_ = -9.01, and log EC_50,*β*Arr2_ = -7.26 corresponding to *Κ*_*β*Arr_ = 5.2 in [42] and log *K*_d_ = -5.70, log EC_50,Gprt,Gs_ = -7.00, and log EC_50,*β*Arr2_ = -6.75 corresponding to *Κ*_*β*Arr_ = 11.2 in [16]. For angiotensin II at the angiotensin II type 1 receptor (AT1R): log *K*_d_ = -7.90, log EC_50,Gprt,IP1_ = -8.84, and log EC_50,*β*Arr2_ = - 7.90 corresponding to *Κ*_*β*Arr_ = 1.0 in [42] and log *K*_d_ = -7.61, log EC_50,Gprt,Gq_ = -8.62, and log EC_50,*β*Arr2_ = - 8.43 corresponding to *Κ*_*β*Arr_ = 6.6 in [53]. One difference is that contrary to most GPCRs, where bound ligands and especially agonist ligands are deeply buried within the receptor [25], the ligands within the binding pocket of MOPr are more exposed to the extracellular surface [59, 60] – see Figure 7 versus Figure 8. This is a likely reason why even potent opioids are rapidly dissociating from their receptor with half-lives of only minutes [59], e.g., 0.5 and 0.7 min for morphine and DAMGO [29], the compounds discussed here, compared to, for example, tiotropium, which has a dissociation half-life (*t*_1/2_ = ln2/*k*_off_ = ln2×*t*_res_) of >30 h at the muscarinic M3 receptor [61, 62]. The resulting short life of the agonist bound MOPr complex could be a possible reason why the concentration-response of arrestin recruitment lags behind the occupancy (is right-shifted) for this pathway.

**Figure 7.**
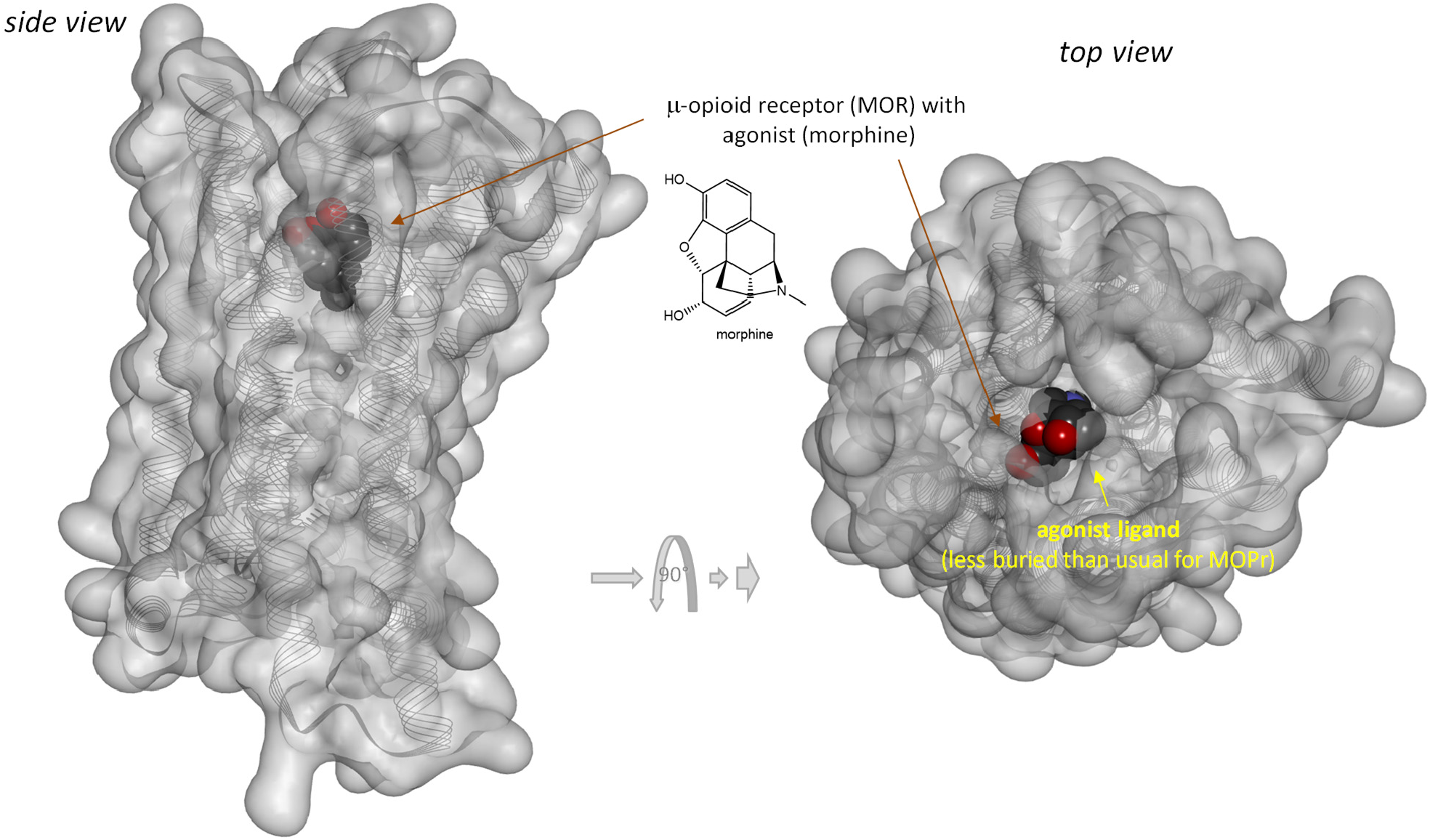
Three-dimensional structure of the classic agonist (morphine) bound form of the *μ*-opioid receptor (MOPr, a type A GPCR). Structure PDB ID# 8EF6 [60] shown covered with a semi-transparent gray surface and the secondary protein-structure indicated; ligand is highlighted as a darker solid CPK structure. Structure is shown from two different perspectives with the one on the right being a 90° rotated and slightly enlarged view from the top. Parts of the ligand are faded as they buried inside the receptor and are obscured by the covering surfaces; however, when looking from the top, part of its surface is not covered and accessible from outside as indicated by its more vivid colors where directly visible.

**Figure 8.**
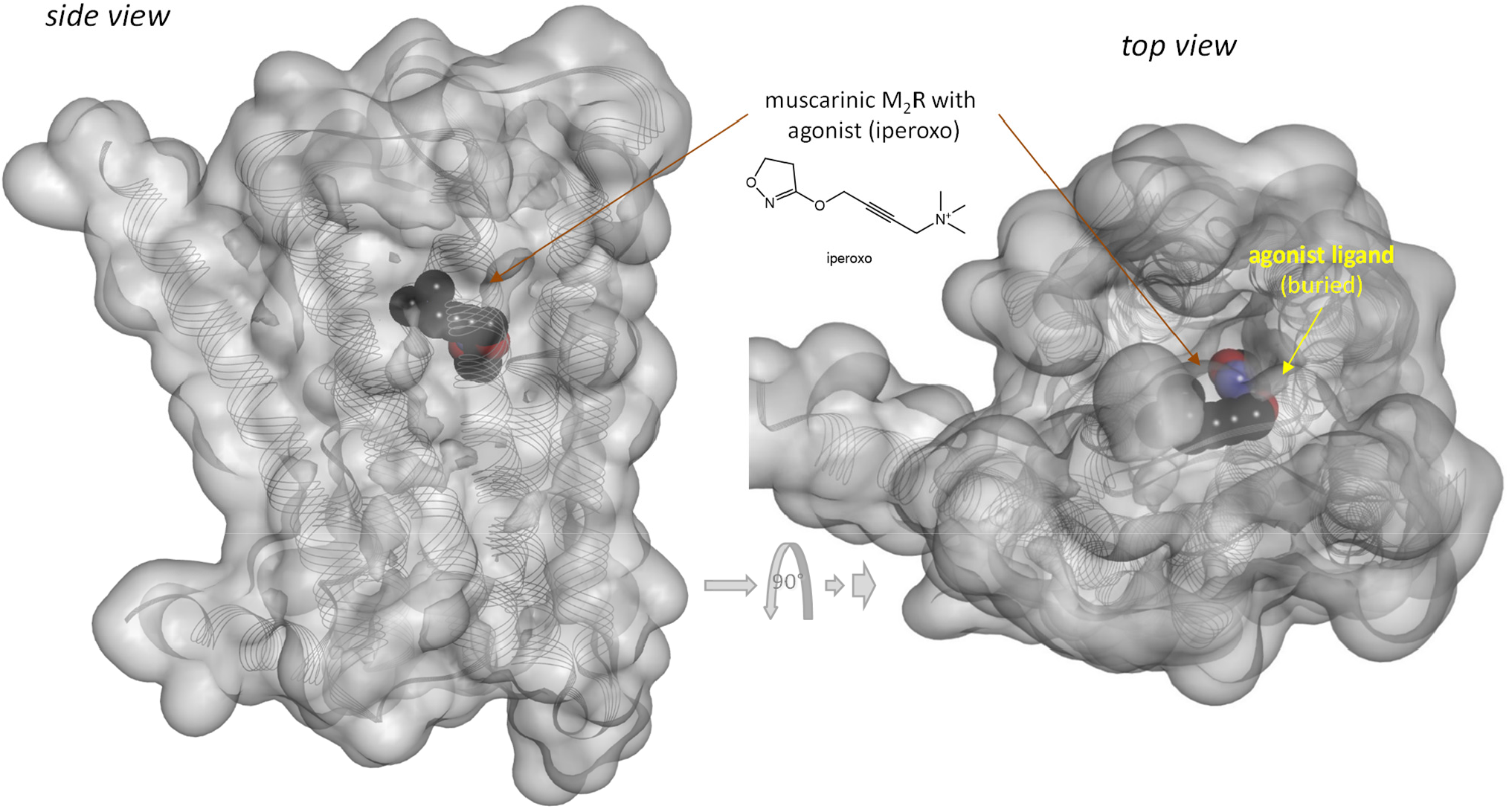
Three-dimensional structure of an agonist (iperoxo) bound form of the muscarinic M2 receptor (a type A*α* GPCR). Structure PDB ID# 4MQS [64] shown covered with a semi-transparent gray surface and the ligand highlighted as a darker solid CPK structure as in the previous figure. Here, the ligand is faded as it is entirely buried inside the receptor and obscured by the covering surfaces from all directions.

As their name implies, a main function of these proteins is to terminate (“arrest”) signaling through GPCRs [63]. For the two known *β*-arrestins (*β*-arrestin-1 and 2, *β*arr1 and *β*arr2 – also known as arrestin-2 and -3, respectively), activation involves two steps: phosphorylation of the activated receptor by specialized GRKs followed by binding of the arrestin(s) to the active phosphorylated receptor to interfere with receptor/G protein coupling [63]. It is conceivable that an “arresting” response is only triggered when a minimum threshold of activation or sufficiently prolonged activation is achieved, and this might require higher agonists concentrations than just binding (occupancy), especially if activated receptors are short-lived. Along similar lines, it is also possible that termination signaling is not triggered unless multiple receptors are activated within the same system, so that low occupancy does not cause response in this pathway. These assumptions, however, are likely to cause responses that are more abrupt than typical hyperbolic ones, which does not seem to be the case here (Figure 3). For example, if two independent receptors within the same system need to be occupied to trigger the response, then *f*_resp_ *∝* (*f*_occup_)^*ν*^ with *ν* = 2, and this produces a response corresponding to *γ* ≈ 0.4 that is just a bit more abrupt than hyperbolic (Supplementary Information, Figure S1); however, larger *ν*s needed for larger shifts result in gradually more abrupt responses.

## Conclusion

To fully connect ligand concentration, receptor occupancy, and assayed responses even in complex cases, models need to include some parametrization to characterize, at a minimum, (1) the ligands ability to bind the receptor (affinity) and (2) activate the bound receptor (efficacy), (3) the degree of activation of unoccupied receptors (efficacy of constitutive activity), (4) the signal modulation along the pathway (gain or loss), and (5) the steepness of concentration dependence (Hill slope). SABRE provides a quantitative model for this that incorporates five parameters in its full form (namely, *K*_d_, *ε, ε*_R0_, *γ*, and *n*) but can be consecutively simplified as needed all the way down to the commonly used Hill or Clark equations by constraining its parameters to specific values. Further, as shown here, it can also be used to fit responses that are not left-but right-shifted compared to occupancy (*Κ* = *K*_d_/EC_50_ < 1) by simply allowing its gain parameter to be less than one indicating an apparent signal attenuation/loss (*γ* < 1). Assays with MOPr provide examples of such data with one left- and one right-shifted response (G protein activation and *β*-arrestin2 recruitment with *Κ*_Gprt_ > 1 and *Κβ*Arr < 1, respectively). As illustrated by fitting such data with SABRE including for oliceridine, weak partial agonists can produce very weak or no activation in the right-shifted pathway without having to be biased agonists due to the combination of low ligand efficacy and signal attenuation.

## Supplementary Information

Supplementary information includes Table S1 and S2 with detailed parameters from fittings shown in Figure 2 and Figure 6, respectively and Figure S1 illustrating a right-shifted response that needs *ν* = 2 independent receptors occupied to trigger it.

## Author Contributions

PB is the sole author; he developed the concept, performed the calculations and data fittings, and wrote the manuscript.

## Acknowledgment

N/A

## Abbreviations

AICc: corrected Akaike information content
DAMGO: D-Ala^2^, N-MePhe^4^, Gly-ol^5^–enkephalin
GPCR: G-protein coupled receptor
GRK: GPCR kinase
MOPr: *μ*-opioid receptor
SABRE: present model (with parameters for Signal Amplification, Binding affinity, and Receptor activation Efficacy)

## Data Availability Statement

Data used for illustrations of model fit are either simulated data generated as described or reproduced from previous publications as indicated in the corresponding figures. The datasets generated and/or analyzed are available from the corresponding author upon reasonable requests.

