## Supplemental Information (Tables S1-S2, Figure S1) for "Quantitative Receptor Model for Responses That Are Left- or Right-Shifted Versus Occupancy (Are More or Less Concentration Sensitive): The SABRE Approach"

*Peter Buchwald*[*]

Department of Molecular and Cellular Pharmacology and Diabetes Research Institute, Miller School of Medicine, University of Miami, Miami, FL, USA

- Supplementary Tables

  Table S1. Detailed parameters from fitting shown in Figure 2.

  Table S2. Detailed parameters from fitting shown in Figure 6.

- Supplementary Figures

  Figure S1. Illustration of a right-shifted response that needs $\nu = 2$ independent receptors occupied to trigger it.

### Supplementary Tables

**Supplementary Table S1**. Parameters and quality of fit descriptors for data shown in Figure 2 (experimental data from [33]). All fittings done in GraphPad Prism.

| Parameter | Phenyl-ephrine | Oxy-metazoline | Naphazoline | Clonidine | Tolazoline | Tena-phtoxaline | Tetra-hydrozoline |
|---|---|---|---|---|---|---|---|
| **A**. Experimental data (from [33])[a] | | | | | | | |
| log $K_d$ | -6.46 | -6.36 | -8.23 | -7.66 | -6.69 | -7.53 | -7.26 |
| log $EC_{50}$ | -7.55 | -6.77 | -8.20 | -7.60 | -6.64 | -7.51 | -7.16 |
| $E_{max,L}$ ($f_{resp,max}$) | 1.000 | 0.730 | 0.480 | 0.330 | 0.100 | 0.270 | 0.070 |
| $\kappa$ (fold vs occup.) | 12.30 | 2.57 | 0.94 | 0.88 | 0.90 | 0.95 | 0.79 |
| **B**. Fit with standard $E_{max}$ (eq. 2) | | | | | | | |
| log $EC_{50}$ | -7.55 | -6.77 | -8.20 | -7.60 | -6.64 | -7.45 | -7.16 |
| $e_{max}$ | 0.996 | 0.729 | 0.480 | 0.332 | 0.100 | 0.276 | 0.068 |
| $r^2$ | 0.996 | 0.998 | 0.997 | 0.994 | 0.949 | 0.977 | 0.978 |
| SSE | 56.25 | 10.50 | 4.26 | 7.52 | 2.95 | 9.41 | 0.33 |
| **C**. Fit with SABRE (eq. 4 with experimental $K_d$) | | | | | | | |
| log $K_d$ (from exp.) | -6.46 | -6.36 | -8.23 | -7.66 | -6.69 | -7.53 | -7.26 |
| $\gamma$ | 11.63 ± 1.83 | | | | | | |
| $\varepsilon$ | 1.000 | 0.177 | 0.065 | 0.038 | 0.009 | 0.029 | 0.006 |
| ↓[b] | | | | | | | |
| log $EC_{50}$ ($K_{obs}$) | -7.53 | -6.82 | -8.46 | -7.81 | -6.73 | -7.65 | -7.29 |
| $e_{max}$ ($f_{resp,max}$) | 1.000 | 0.715 | 0.447 | 0.313 | 0.096 | 0.261 | 0.065 |
| $\kappa$ | 11.63 | 2.88 | 1.69 | 1.40 | 1.10 | 1.31 | 1.06 |
| $r^2$ | 0.996 | 0.997 | 0.951 | 0.974 | 0.943 | 0.941 | 0.959 |
| SSE | 58.54 | 16.88 | 75.88 | 31.64 | 3.304 | 24.27 | 0.65 |
| **D**. Fit with SABRE for $f_{resp}$ vs. $f_{occup}$ directly (eq. 17) | | | | | | | |
| $\gamma$ | 11.63 ± 1.83 | | | | | | |
| $\varepsilon$ | 1.000 | 0.177 | 0.065 | 0.038 | 0.009 | 0.029 | 0.006 |
| $r^2$ | 0.996 | 0.997 | 0.951 | 0.974 | 0.943 | 0.941 | 0.959 |
| SSE | 58.54 | 16.88 | 75.88 | 31.64 | 3.304 | 24.27 | 0.65 |

[a] Experimental data – average of log $K_A$ and log $K_B$ from Table 3 and 4 in [33]; $\kappa$ (fold shifts vs occupancy) calculated from the $K_d$/$EC_{50}$ values. [b] Derived values for the present model (using eqs. 6, 7, and 17). Quality of fit descriptors included: $r^2$, correlation coefficient; SSE, sum of squared errors.

**Supplementary Table S2**. Parameters and quality of fit descriptors for data shown in Figure 6 (experimental data from [29]). All fittings done in GraphPad Prism.

| Parameter | Loperamide | DAMGO | Morphine | Buprenorphine | Oliceridine (R)-TRV130 | (S)-TRV130 |
|---|---|---|---|---|---|---|
| **A**. Experimental data (from [29]) | | | | | | |
| log $K_d$[a] | -7.64 | -7.34 | -7.02 | -8.37 | -7.29 | -5.73 |
| log $EC_{50,Gprt}$ | -9.20 | -8.61 | -8.17 | -9.26 | -8.49 | -6.53 |
| $E_{max,Gprt}$ ($f_{resp,max}$, %) | 98 | 99 | 98 | 84 | 82 | 85 |
| log $ED_{50,\beta Arr}$ | -6.94 | -6.10 | -6.00 | | | |
| $E_{max,\beta Arr}$ ($f_{resp,max}$, %) | 51 | 99 | 25 | | | |
| **B**. Fit with standard $E_{max}$ (eq. 2) | | | | | | |
| log $EC_{50,\ Gprt}$ | -8.98 | -8.62 | -8.09 | -8.66 | -8.51 | -6.53 |
| $e_{max,Gprt}$ | 105.4 | 96.9 | 98.1 | 83.8 | 84.3 | 86.6 |
| log $EC_{50,\beta Arr}$ | -6.89 | -6.12 | -6.10 | | | |
| $e_{max,\beta Arr}$ | 52.9 | 97.9 | 23.1 | | | |
| **C**. Fit with SABRE (eq. 4 with experimental $K_d$) | | | | | | |
| log $K_d$ (from exp.)[a] | -7.64 | -7.34 | -7.02 | -8.37 | -7.29 | -5.73 |
| $\gamma_{Gprt}$ | 19.88 | | | | | |
| $\gamma_{\beta Arr}$ | 0.067 | | | | | |
| $\varepsilon$ | 0.950 | 0.998 | 0.792 | 0.141 | 0.382 | 0.258 |
| ↓[b] | | | | | | |
| log $EC_{50,\ Gprt}$ | -8.98 | -8.70 | -8.29 | -8.98 | -8.26 | -6.55 |
| $e_{max,Gprt}$ | 99.8 | 100.0 | 98.9 | 79.1 | 93.4 | 88.9 |
| log $EC_{50,\beta Arr}$ | -6.69 | -6.17 | -6.43 | -8.31 | -7.10 | -5.61 |
| $e_{max,\beta Arr}$ | 55.1 | 96.7 | 19.9 | 1.1 | 3.9 | 2.2 |

[a] Experimental data from [29]. [b] Derived values for the present model (using eqs. 6 and 7).

### Supplementary Figures

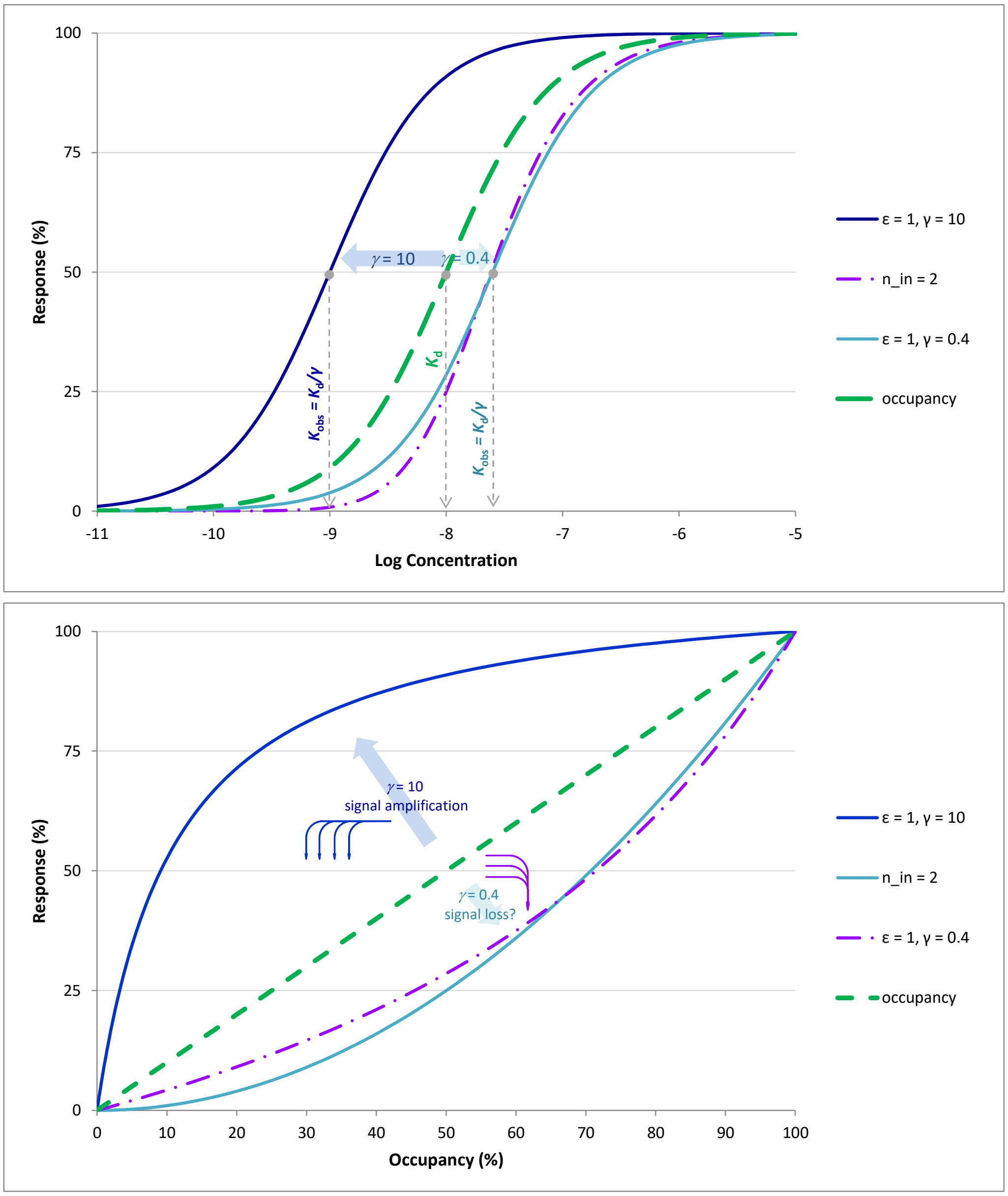


**Figure S1**. Illustration of a right-shifted response (dashed purple line) caused by the assumption that $\nu = 2$ independent receptors within the same system need to be occupied to trigger the response, which result in $f_{resp} \propto (f_{occup})^{\nu}$. Responses are shown as a function of log concentration (top) or fractional occupancy (bottom).
